# Excitatory Dysfunction and Phenotypic Rescue in a Human Neuronal Model of *SCN2A*-Related Disorders

**DOI:** 10.64898/2026.05.18.725745

**Authors:** Sashmita Panda, Diana Jazmin Ixmatlahua Ribera, Luis A. Williams, Sheng Tang, Karthiayani Harikrishnan, Vaibhav Joshi, Steven J. Ryan, Chinwendu Obi Obasi, Linda Laux, Enrique Rojas, Owen McManus, Graham T. Dempsey, Alfred L. George, Scott K. Adney

## Abstract

*SCN2A*-related disorders result from pathogenic variants in the gene encoding for the voltage-gated sodium channel Nav1.2. Collectively, these disorders result in variable age of onset epilepsy, autism spectrum disorder, and epileptic encephalopathies. While the mechanisms of haploinsufficiency resulting in autism spectrum disorder have been explored in detail, few studies report the impact of pathogenic missense variants in human neurons. In this work, we combined conventional electrophysiology and high-throughput all-optical electrophysiology assays to analyze the *SCN2A* p.M1879T pathogenic variant associated with early-onset epilepsy and developmental delay. In both platforms, iPSC-derived excitatory neurons expressing the disease variant showed greater firing at higher stimuli compared to the isogenic control neurons (corrected by CRISPR/Cas9), as well as changes to action potential shape (steeper slope and larger amplitude) with evoked firing. We used machine learning techniques on the optical physiology dataset to classify the two genotypes, finding that sodium channel blocking anti-seizure drugs could restore an isogenic phenotype. This work demonstrates proof of sodium channel blocker efficacy in a human neuronal model of *SCN2A*-related epilepsy and highlights the power of leveraging high-throughput all-optical electrophysiology for testing drug efficacy.

## Introduction

Variants in the gene *SCN2A*, which encodes for the Nav1.2 voltage-gated sodium channel, are associated with monogenic causes of variable age of onset epilepsy, autism spectrum disorder, and epileptic encephalopathies^1^. Collectively, these are termed *SCN2A*-related disorders, and have variable phenotypic severity, ranging from a self-limited syndrome of neonatal seizures, often familial^2–4^, to severe epileptic encephalopathy with significant neurodevelopmental impairment^5,6^. *SCN2A* is also a high-risk autism gene and its haploinsufficiency is causative of autism spectrum disorder with and without epilepsy. In individuals with autism spectrum disorder and intellectual disability, the *SCN2A* gene has been found to have the highest number of *de novo* variations, primarily truncations resulting in nonfunctional channels^7^.

The Nav1.2 channel encoded by *SCN2A* is highly expressed in excitatory neurons, with high channel concentration especially in the axon initial segment (AIS) early in development. There is subsequently a developmental switch where Nav1.6, encoded by *SCN8A*, expresses throughout the AIS with Nav1.2 restricted to the proximal segment of the AIS^8^. Due in part to its high representation in autism, the circuit and behavioral mechanisms of *SCN2A* haploinsufficiency have been explored in detail^9–15^. In contrast, less is known about the impact of missense *SCN2A* variants in human pathophysiology due to a paucity of studies looking at neuronal function^16^, especially in patient-specific human neuronal model systems^17–19^.

Early work in heterologous cells examined the functional properties of pathogenic variants in *SCN2A*, concluding that variants associated with epileptic encephalopathies were primarily gain-of-function while late-onset seizures and autism were due to loss-of-function variants^20,21^. Recent work has shown that this genotype-phenotype correlation does not always hold. High-throughput patch-clamp analysis of variants in *SCN2A*-related disorders has shown complex biophysical properties, including mixed properties^22,23^.

To extend insights gained from ion channel biophysics to functional neuron activity, *in silico* modeling has been applied^24^. These models, while promising, often lack validation in experimental model systems. In this work, we utilized neurons derived from patient-induced pluripotent stem cells (iPSCs) to examine the impact of a gain-of-function *SCN2A* variant, p.M1879T, predominantly affecting channel inactivation^22,25^. We applied high-throughput all-optical electrophysiology (*Optopatch*) measurements^26,27^ and compared the *SCN2A* p.M1879T patient-derived neurons with CRISPR/Cas9-corrected isogenic control neurons. We validated and extended the findings from these ‘*Optopatch*’ experiments with traditional patch-clamp electrophysiology. Furthermore, we evaluated in *SCN2A* p.M1879T neurons the effect of traditional and newer sodium channel blocking compounds on evoked neuronal firing, demonstrating the translational potential of high-throughput functional measurements and analyses.

## Results

### Generation of patient-derived p.M1879T excitatory neurons

An iPSC line was generated from a male child with early-onset epilepsy heterozygous for the variant *SCN2A*-p.M1879T (c.5636T>C), initially classified as a variant of uncertain significance (VUS) with a more recent status of pathogenic/likely pathogenic (P/LP). This *SCN2A* variant has been reported in at least three unrelated individuals in ClinVar (NCBI). The child presented with mild global developmental delay and infantile-onset tonic seizures that were initially unresponsive to anti-seizure medication trials^25^. Following administration of the sodium channel blocker phenytoin, the patient’s seizures resolved. He was then transitioned to carbamazepine monotherapy. On follow-up at 10 years old, the patient is nonverbal and has been diagnosed with autism spectrum disorder. He is reported to have a few short focal seizures per year with no convulsions. He remains on carbamazepine monotherapy.

The *SCN2A* gene product Nav1.2 is highly expressed throughout the postnatal brain with high concentration in cortical excitatory neurons^21^. To examine the neuron dysfunction of patient-derived p.M1879T excitatory neurons, we differentiated the patient iPSCs into cortical excitatory neurons using transcriptional programming mediated by overexpression of NEUROG2 (NGN2) combined with patterning by small molecule inhibition of SMAD and Wnt signaling pathways, as detailed in methods. These differentiated NGN2-induced neurons (iNeurons) were cultured on rodent glia to accelerate functional maturity and produce homogeneous electrophysiological properties^28,29^.

For a comparison control group, we generated isogenic control iPSC lines with CRISPR/Cas9 editing and confirmed correction of the *SCN2A* pathogenic mutation with Sanger sequencing. The parental p.M1879T iPSC line and its respective controls were confirmed to have a normal 46XY karyotype (**Supplementary Figure 1A**) and no potential off-target editing was detected at the *SCN1A* or *SCN3A* orthologous sites. Additional isogenic iPSC clones were also obtained that went through the editing process but did not correct the initial *SCN2A* pathogenic mutation. Neurons differentiated from these ‘processed controls’ clones were used in the all-optical electrophysiology experiments and were designated as ‘p.M1879T lines’. Differentiated neurons expressed the pan-neuronal marker MAP2 (Microtubule-Associated Protein 2) and the expected excitatory glutamatergic marker vGlut1, as determined by immunocytochemistry (**Supplementary Figure 1B**). There was no difference seen in total *SCN2A* transcript expression in neuronal lines using digital droplet PCR (ddPCR). An overview of the experimental design is shown in **Figure 1**.

**Figure 1.**
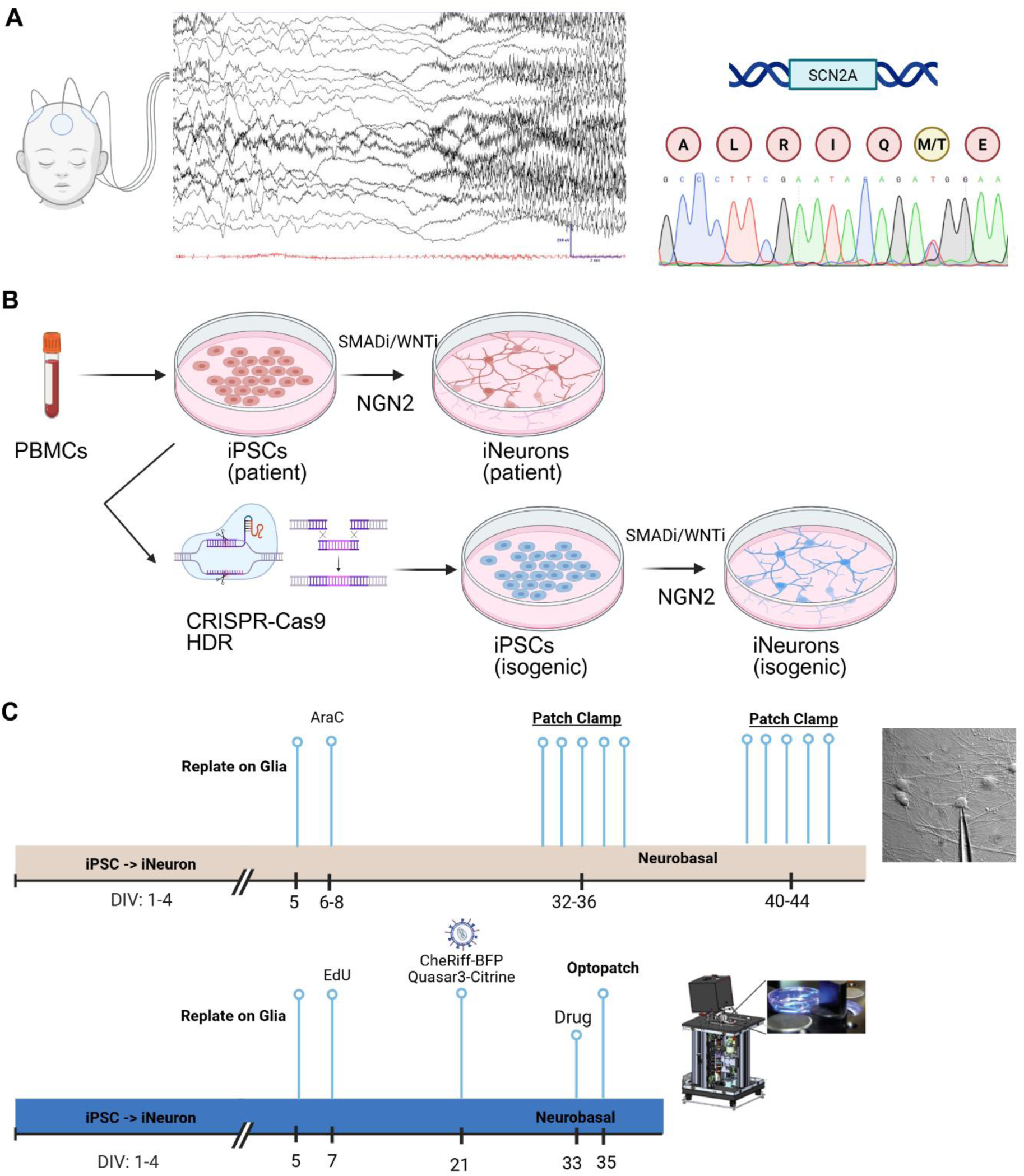
Overview of Analysis of p.M1879T patient-derived Neurons A. *Left*, EEG showing tonic seizure of the patient heterozygous for the p.M1879T missense variant (*right*) in *SCN2A*. B. Schematic of iPSC creation, CRSPR/Cas9 editing, and neuron differentiation using dual SMAD/Wnt pathway inhibition combined with NGN2 overexpression. C. Timeline of neuron culture and differentiation/maturation process for path-clamp electrophysiology (*top*) and all-optical ‘Optopatch’ electrophysiology (*bottom*), with experimental DIV timepoints shown below.

### Aberrant Neuronal Firing in p.M1879T Neurons

To characterize the *SCN2A*-p.M1879T patient-derived cortical excitatory neurons, we employed whole cell patch-clamp electrophysiology to compare the functional properties of p.M1879T neurons to an isogenic control at two different time points, day *in vitro* (DIV) 32-36 and DIV 40-44. First, we analyzed the intrinsic excitability of the two *SCN2A* genotypes by recording action potentials (APs) in response to increasing the current injection. At the earlier culture timepoint (DIV 32-36), when we compared the number of APs in response to current injections (**Figure 2A**), we found that the p.M1879T neurons fired at higher frequencies with higher current stimuli compared to the isogenic control neurons (p.M1879T: 15.0 ± 2.3 vs isogenic: 7.8 ± 1.5 spikes at 100 pA, p<0.05, Mann-Whitney). Analysis of the waveform of the first AP (**Figure 2B**) showed that the p.M1879T neurons exhibited greater peak amplitude (p.M1879T: 63.5 ± 1.7 mV vs isogenic: 55.3 ±1.5 mV, p<0.05 by Mann-Whitney) and larger maximal rise slope (dV/dt) (p.M1879T: 70.1 ± 5.2 mV/ms vs isogenic: 53.8 ± 3.2 mV/ms, p<0.05 by Mann-Whitney) (**Figure 2C**). Other parameters including rheobase, action potential threshold, time to max rise slope, rise time constant and area were not different between the two *SCN2A* genotypes (**Supplementary figure 2 A**).

**Figure 2.**
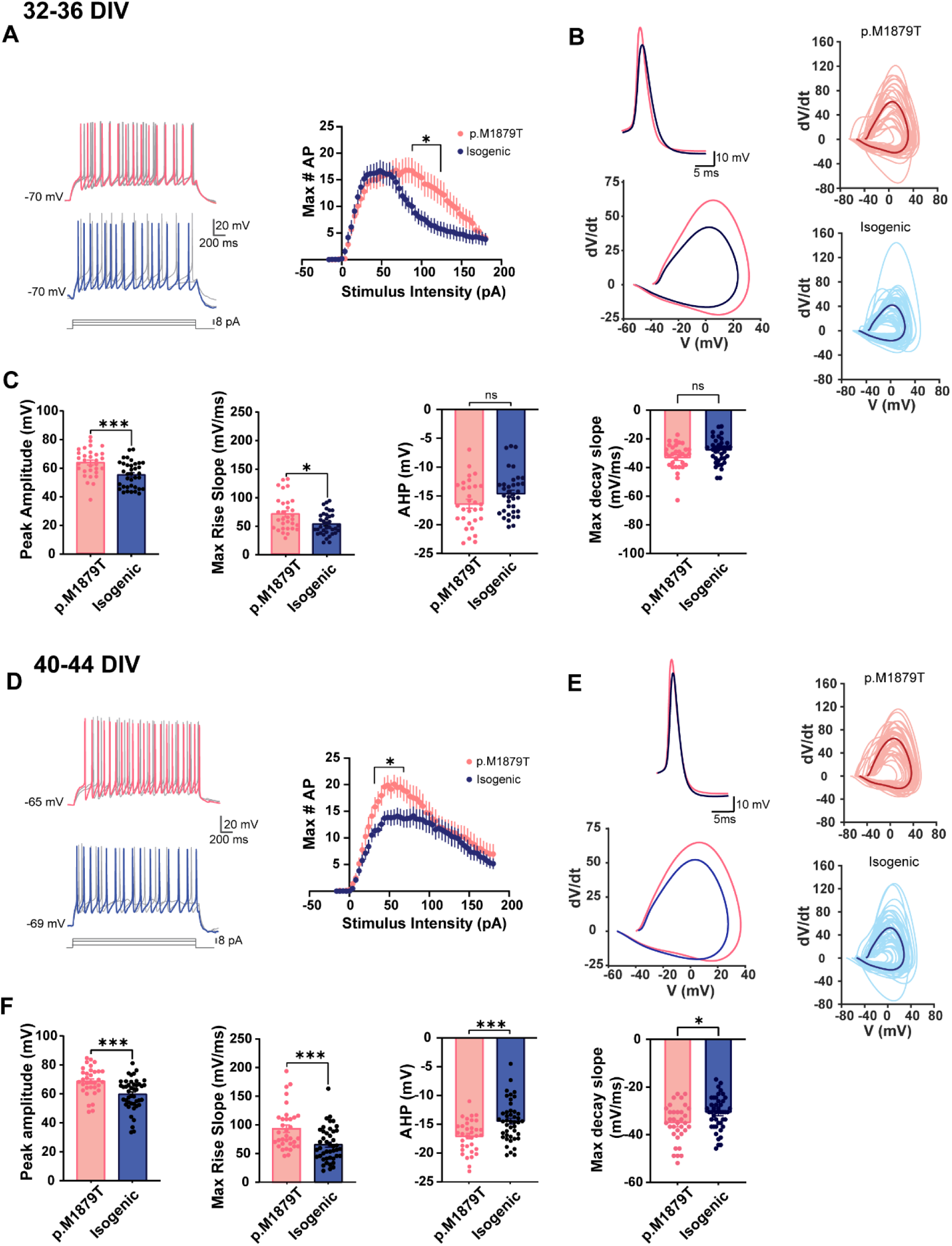
Electrophysiological profile of p.M1879T Neurons. A, D. *Left:* Representative traces of current clamp recordings in neurons showing action potentials generated by current injection (32-40pA, 2000 ms). p.M1879T neurons (red) and isogenic neurons (blue) from DIV32-36 (**A**) and DIV40-44 (**D**). *Right*: Summary of the number of APs evoked by step depolarizing current injections (−16 to 180 pA); DIV32-36: n=31 neurons for p.M1879T vs n=35 neurons for isogenic; DIV40-44: n=34 neurons for p.M1879T vs n=44 neurons for isogenic. *, p<0.01, Mann-Whitney with Dunn’s post-hoc test. B, E. *Left up*: Inset of average AP in a p.M1879T neuron (red) and in an isogenic neuron (blue). B, E: Left down: Average Phase plots (d*V*/d*t* versus V) of the APs from top. B, E right. Phase plot of all the neurons, traces in light color are the phase plot of individual cells, the traces in dark color represent the average phase plot for each group: DIV32-36 (**B**) and DIV40-44 (**E**). C, F. Plots showing action potential properties from left to right: peak amplitude (mV), Max rise slope (d*V*/d*t*), AHP (mV) and Max decay slope at DIV32-36 (**C**) and DIV40-44 (**F**). *p<0.05 by Mann-Whitney; *p<0.01 by Mann-Whitney. Data are shown as mean ± SEM.

At the later timepoint (DIV 40-44), the difference in intrinsic excitability was evident at lower current injections with neurons expressing the *SCN2A* pathogenic variant demonstrating a higher number of APs (p.M1879T: 20.1 ± 1.8 spikes vs isogenic: 13.9 ± 1.5 spikes at 50 pA current stimulation, p<0.05, Mann-Whitney). The stimulus-firing curves are shown in **Figure 2D**. Similar to the previous timepoint, both the amplitude of the action potential and maximal slope (dV/dt) were significantly higher in the p.M1879T neurons compared to the isogenic control (peak amplitude, p.M1879T: 68.6 ± 1.6 mV vs isogenic: 59.7 ± 1.6 mV, p<0.05 by Mann-Whitney; Max rise slope, p.M1879T: 93.1 ± 6.3 mV/ms vs isogenic: 65.4 ± 4.5 mV/ms, p<0.05 by Mann-Whitney) (**Figure 2E** and **Figure 2F**). Unlike the previous timepoint, the fast AHP was significantly larger (deeper afterhyperpolarization) and the AP decay had higher max velocity for the p.M1879T neurons (fAHP, p.M1879T: −17.08 ± 0.47 mV vs isogenic: −14.43 ± 0.5 mV, p<0.05, Mann-Whitney; AP max decay slope p.M1879T: −35.3 mV/ms ± 1.3 vs isogenic: −30.95 ± 1.04 mV/ms, p=0.0124, Mann-Whitney). As in DIV 32-36, no significant changes were detected in other functional parameters examined (**Supplemental Figure 2B)**. Taken together, these results demonstrate how the biophysical changes in p.M1879T channels result in hyperexcitable properties in patient-derived neurons.

### All-Optical Electrophysiology of p.M1879T Neurons

In addition to the low-throughput methodology of whole-cell patch-clamp recordings, we conducted our functional analysis with all-optical electrophysiology, which further allowed for pharmacological profiling. To this end, two *SCN2A* neuronal lines heterozygous for the p.M1879T variant (called ‘p.M1879T lines [+/M1879T]) were cultured and characterized alongside the isogenic *SCN2A* corrected control lines (termed ‘Isogenic lines [+/+]’) on the all-optical electrophysiology ‘Optopatch’ platform^26,27^ at DIV 35 (**Figure 1C**). All these neuronal lines were derived from the patient founder *SCN2A* p.M1879T iPSC line and went through the same CRISPR/Cas9 treatment and single-cell cloning steps, with the two ‘p.M1879T [+/M1879T]’ lines retaining the *SCN2A* pathogenic genotype and the two ‘Isogenic [+/+]’ lines showing genetic correction of the variant. Optogenetic activation protocols were run to sequentially excite the neurons, as shown in **Figure 3A**. For analysis, we applied two thresholds; we selected neurons with an average signal to noise ratio (SNR) at least 6, and we included neurons with at least two identified spikes throughout the entire protocol. In total, 1,192,149 spiking events were analyzed in 33,440 iPSC-derived neurons.

**Figure 3.**
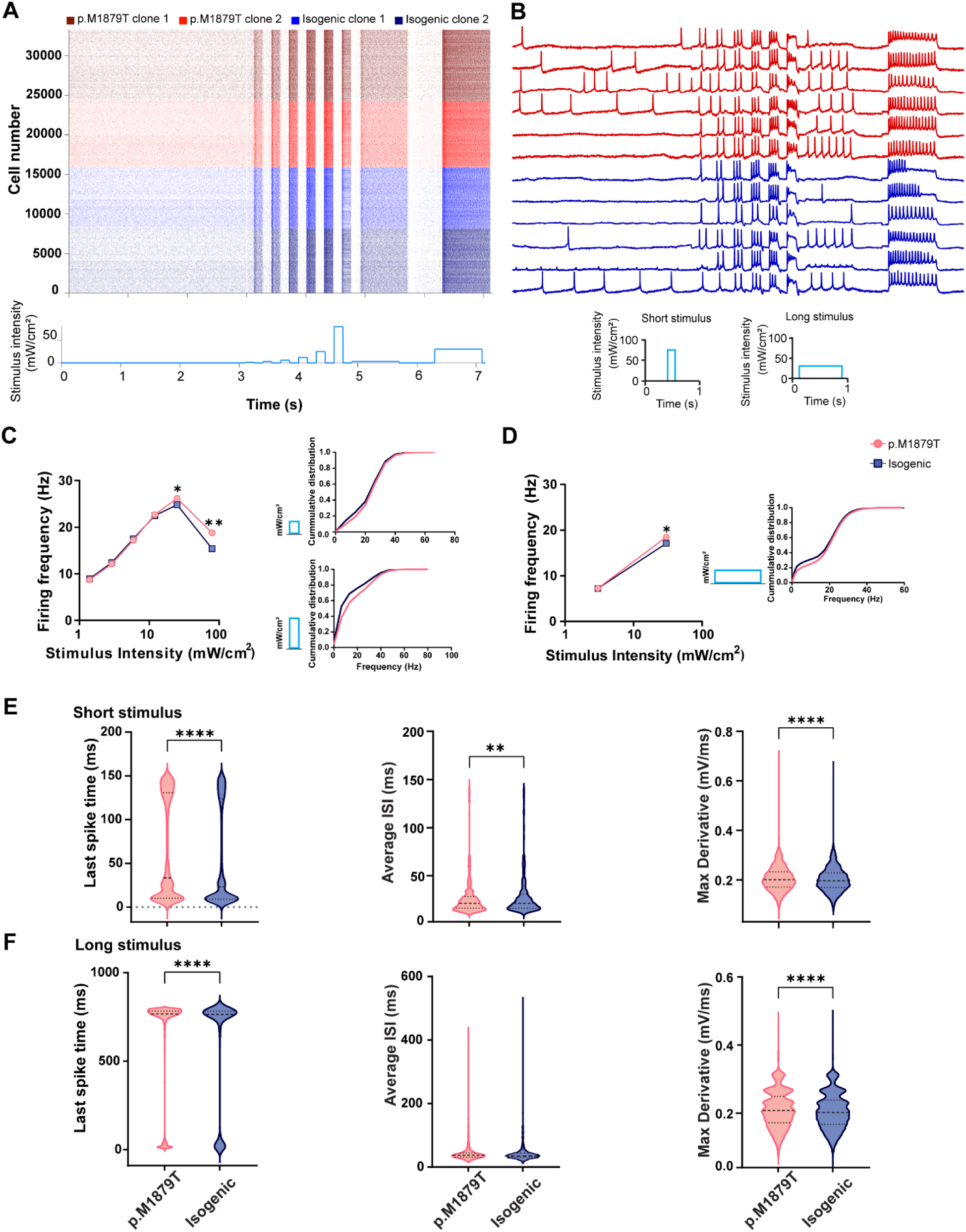
Optopatch Phenotyping of p.M1879T Neurons Shows Phenotypic Separation at Higher and Longer Stimuli A. Raster plot of neuronal firing (top) and stimulus protocol (bottom) from one optopatch run; B. Representative neuron optical voltage traces from patient lines (red) and isogenic control lines (blue). C. *Left*: Stimulus-frequency plot of p.M1879T (red circles) vs isogenic (blue squares) for short (150ms) optical stimuli showing significant differences at highest stimuli (*,p<0.001 by Kruskal-Wallis with Dunn’s post-hoc test); *right*: Cumulative distribution of firing frequency for neurons at specified optical stimuli. D. Stimulus-frequency plot for longer (800 ms) optical stimuli (*,p<0.001 by Kruskal-Wallis with Dunn’s post-hoc test) E and F. Parameters at highest optical stimuli (79 mW/cm^2^ and 30 mW/cm^2^) from (C and D, respectively) shown: average ISI, last spike time, firing frequency, and maximum derivative; ****p<0.001 by Mann-Whitney test.

We first analyzed the firing frequency vs. stimulus intensity, a surrogate of the current-frequency curve in our conventional current clamp experiments. The optical stimulus protocol includes 6 short duration step stimuli (150 ms) and 2 long duration step stimuli (800 ms; **Figure 3A**, **bottom**). Two replicate ‘runs’ consisted of independently plated neurons transduced with lentiviral particles for *Optopatch* functional analysis. As shown in **Figure 3C**, the average firing frequency of the p.M1879T lines [depicted as ‘+/M1879T’] was significantly increased compared to the isogenic controls [‘+/+’] at the higher stimulus intensities of 25 and 79 mW/cm^2^, with the highest effect size detected at 79 mW/cm^2^. The detailed results from each run and the combined results are displayed in **Table 1**. Similarly, at the highest intensity of the longer duration steps (30 mW/cm^2^), the ‘+/M1879T’ lines displayed higher firing frequency compared to the isogenic ‘+/+’ lines, but there was no significant difference observed at the lower stimulus intensity, **Figure 3D**. These differences persisted when single-spiking neurons were excluded, confirming robustness of the result. Since the p.M1879T neurons fire at a higher rate than the isogenic control neurons, we also assessed the timing of the last evoked spike in response to the highest stimuli (79 mW/cm^2^). Indeed, the p.M1879T lines continued to fire significantly later into the stimulus (**Figure 3E and 3F**), indicating less susceptibility to depolarization block compared to the isogenic lines. Additionally, the average interspike interval (ISI) was shorter in p.M1879T neurons and the maximum derivative of the optical action potential (a surrogate marker of dV/dt) was significantly greater. Taken together, these findings obtained with the higher throughput optical physiology measurements show that the p.M1879T lines have a reduced ‘failure rate’, firing faster at higher and longer stimulus intensities when compared to isogenic control neurons.

**Table 1:** Key Optopatch Parameters.

| Genotype | Firing Freq at 25 mW/Cm2 (short) | Firing Freq at 79 mW/cm2 (short) | Firing Freq at 3 mW/Cm2 (long) | Firing Freq at 30 mW/cm2 (long) | Number of neurons analyzed (n) |
| --- | --- | --- | --- | --- | --- |
| p.M1879T (run1) | 25.7* +/- 0.2 [d=0.11] | 17.6** +/- 0.3 [d=0.23] | 7.31 +/- 0.1 | 18.2** +/- 0.2 [d=0.12] | 8,475 |
| Isogenics (run1) | 24.5 +/- 0.5 | 14.6 +/- 0.3 | 7.44 +/- 0.1 | 16.9 +/- 0.3 | 7,761 |
| p.M1879T (run2) | 26.8 * +/- 0.3 [d=0.15] | 19.8** +/- 0.3 [d=0.26] | 7.20 +/- 0.09 | 18.9** +/- 0.3 [d=0.13] | 9,058 |
| Isogenics (run2) | 25.1 +/- 0.3 | 16.3 +/- 0.3 | 7.04 +/- 0.1 | 17.4 +/- 0.3 | 8,146 |
| p.M1879T (combined) | 26.2* +/- 0.2 [d=0.13] | 18.8** +/- 0.2 [d=0.25] | 7.25 +/- 0.1 | 18.5** +/- 0.2 [d=0.13] | 17,533 |
| Isogenics (combined) | 24.8 +/- 0.2 | 15.4 +/- 0.2 | 7.24 +/- 0.1 | 17.1 +/- 0.2 | 15,907 |

Firing Frequency parameters for specified Optopatch stimuli. Numbers are Hz +/- 95% CI of the mean value. \*: $p < 0.001$ , Kruskal-Wallis test with Dunn's correction for multiple comparisons; \*\* indicates higher effect size [cohen's D]

### Restoration of p.M1879T functional phenotype with sodium channel blockers

We next turned to a particular benefit of the all-optical electrophysiology approach: the ability to perform high-throughput pharmacology. Using this functional platform, we were able to test the effect of several sodium channel blockers on the p.M1879T and isogenic neuronal lines in parallel. We selected two sodium channel blocking anti-seizure drugs to which the patient had previously responded, phenytoin (PHT) and carbamazepine (CBZ). We also assessed the effect of treatment with newer sodium channel blocking agents, lacosamide (LAC) and cenobamate (CBM), which have distinct mechanisms from the classic use-dependent sodium channel blocking agents^30,31^. These drug treatments were applied over a multi-logarithmic concentration range and compared to the DMSO vehicle control. In all cases, the cultured neurons were exposed to the drugs for 48 hours, and fresh drug or vehicle was added prior to optical physiology measurements as in **Figure 1C**.

In **Figure 4A-D**, the effect of each drug on the stimulus-frequency plot is shown for p.M1879T neurons along with the isogenic (in DMSO vehicle) for comparison. At higher concentrations, all sodium channel blocker drugs display a use-dependent reduction in firing, where the effect was strongest at the higher stimuli. Examining the higher stimulation paradigms in which we detected phenotypic separation between the p.M1879T lines and Isogenic controls, we then compared whether treatments of p.M1879T lines with the different drugs could reverse their firing phenotype, bringing their neuronal properties closer to those of isogenic control neurons (**Figure 4E-H**). All drugs showed efficacy in bringing the firing rate of p.M1879T neurons closer to the mean value of Isogenic neurons. We defined phenotypic rescue if the firing rate was significantly different from the p.M1879T vehicle level but was not significantly different from the isogenic level (i.e., showed overlap). The concentration for each drug achieving this phenotypic rescue was: PHT 30 μM, LAC 300 μM, and CBZ 100 μM (circled in **Figure 4E-H**). CBM did not demonstrate phenotype reversal by these criteria. Higher drug concentrations resulted in significantly lower firing compared to the isogenic lines, consistent with over-correction of the firing phenotype. In each case, drug treatment of the isogenic line resulted in further reduction of evoked firing.

**Figure 4.**
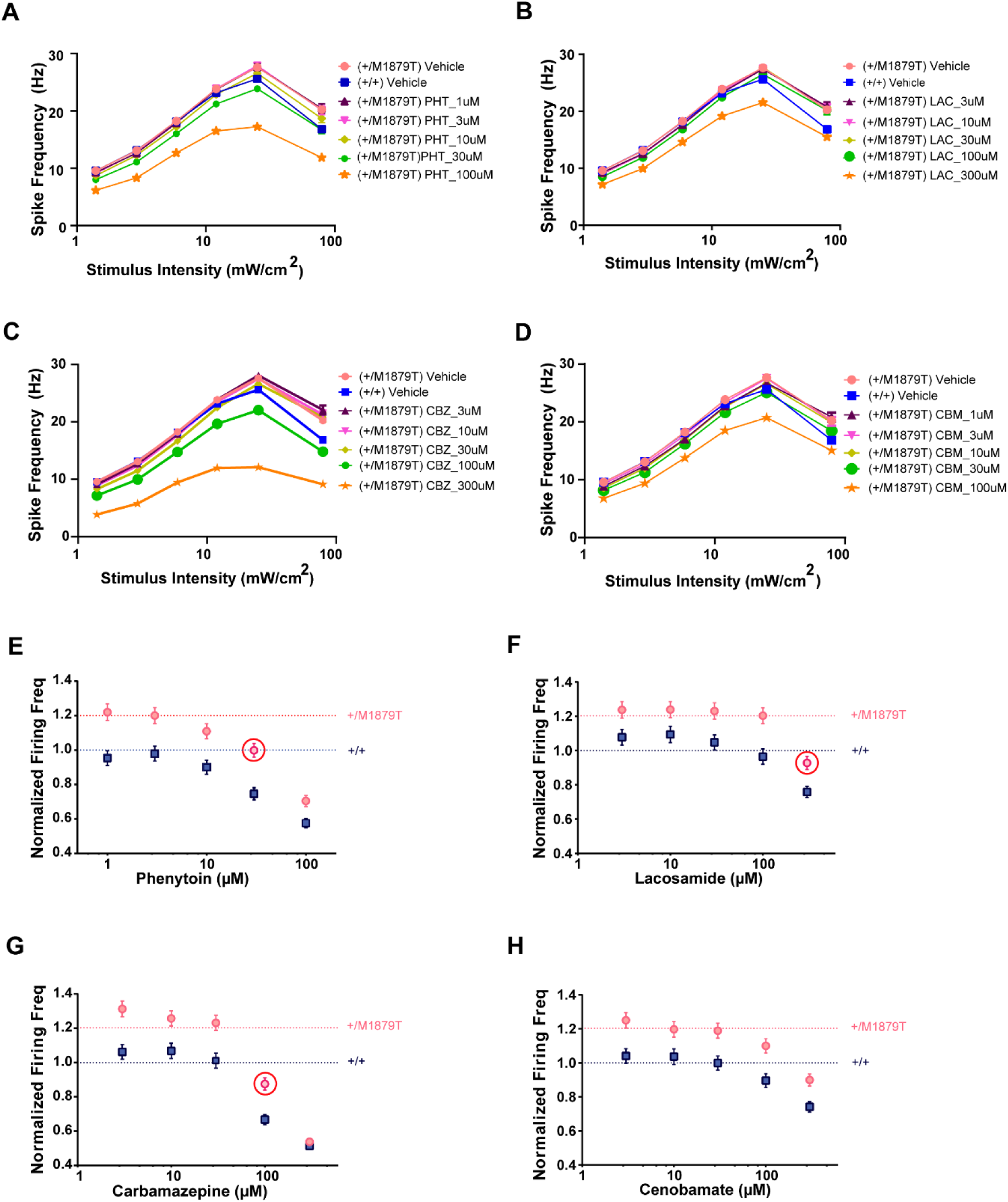
Pharmacology of p.M1879T Neurons Reverses Firing Phenotype A-D. Stimulus-frequency plot of p.M1879T neurons with increasing drug concentrations, with the isogenic neuron properties shown for reference. Phenytoin in **A** (PHT), lacosamide in **B** (LAC), carbamazepine in **C** (CBZ), cenobamate in **D** (CBM). E-H. Normalized firing frequency at the highest stimuli (79 mW/cm^2^) showing dose-dependent reduction of firing of p.M1879T neurons (red circles) and isogenic neurons (blue squares). Values are normalized with respect to the isogenic properties.

To further define the impact of sodium channel blockers on the *SCN2A*-p.M1879T patient-derived neurons, we used a machine learning classification approach to perform unbiased analysis of the evoked firing features. We first took average features from random subpopulations containing 100 neurons from the p.M1879T and Isogenic lines, using bootstrap sampling to reduce variance. We then applied random forest classification with feature selection for training the model on the dataset without exposure to any drugs. The model displayed high accuracy (>90%) differentiating vehicle-treated p.M1879T and isogenic lines using this approach. Furthermore, we found that classification accuracy was reduced in a dose-dependent manner with each sodium channel blocker, consistent with alteration of the evoked firing feature parameter set (**Figure 5** and **Supplementary Figure 3**). In some cases, defined by a classification accuracy below 50%, the drug-treated neurons showed over-correction. In particular, the following concentration ranges were found to be in-between phenotypic reversal and over-correction: PHT 10 μM to 30 μM; CBZ 30 μM to 100 μM; LAC 100 μM to 300 μM; CBM 10 μM to 30 μM. To verify predictions from the model, we looked at aggregate firing features with principal component analysis (PCA) in individual runs (**Figure 5** and **Supplementary Figure 4**). We quantified separation by analyzing the distance from the centroid of each drug projection relative to vehicle treatment. Low and intermediate drug concentrations show overlap with the untreated isogenic lines, consistent with partial correction of the phenotype. Higher concentrations of the applied drugs result in over-correction of the phenotype, as evidenced from the distance map demonstrating large deviation in the feature space. Taken together, these data indicate the ability of sodium channel blocking anti-seizure drugs to restore the aberrant neuronal firing of p.M1879T excitatory neurons, providing *in vitro* validation in human neurons of a practice clinically employed in *SCN2A*-related disorders.

**Figure 5.**
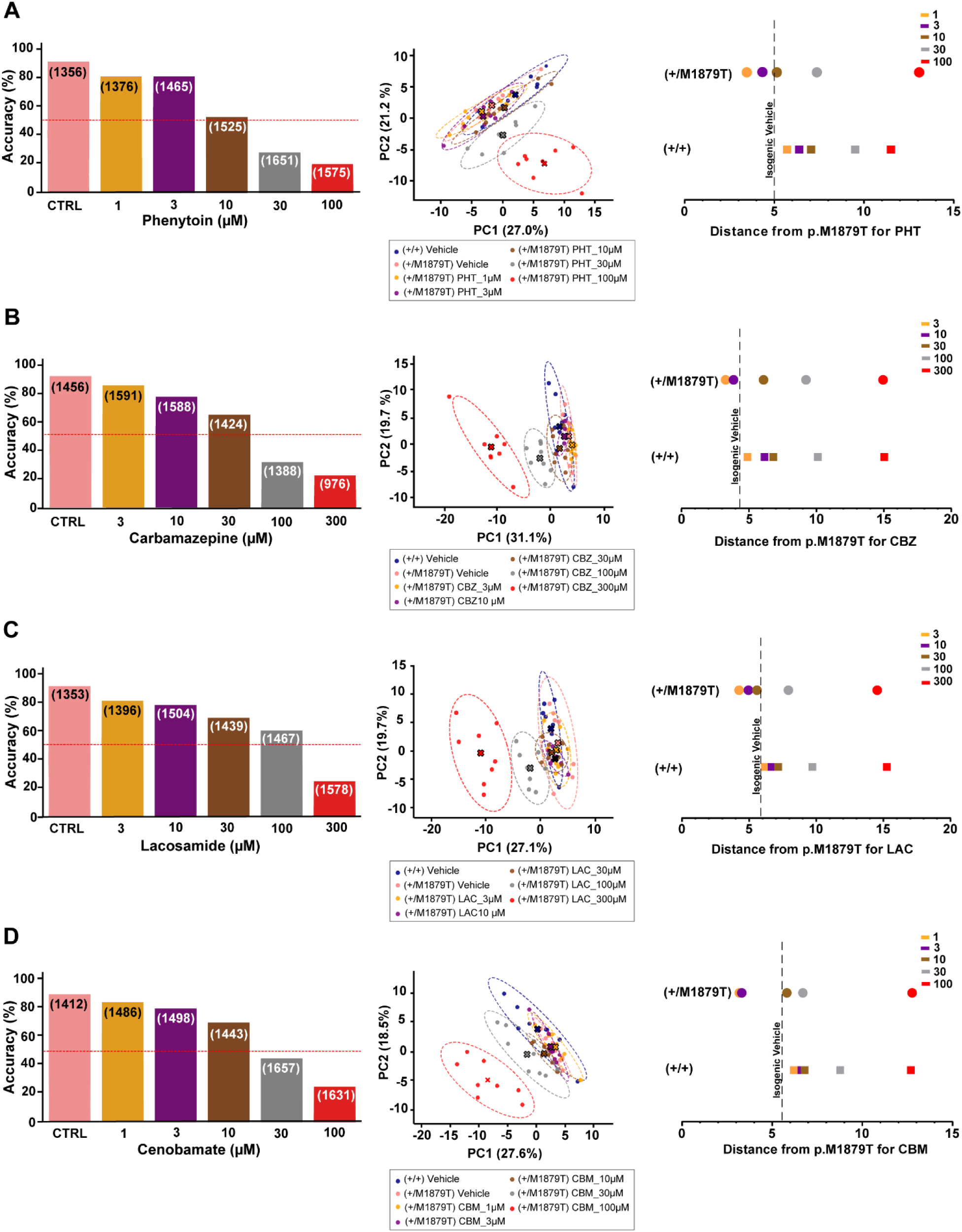
Classification Model Highlights Phenotypic Rescue with Sodium Channel Blockers A-D. *Left:* Classification accuracy with random forest feature selection model showing reduction in classification accuracy with increasing concentrations of indicated drug. *Middle*: PCA plots showing aggregate feature clusters with respect to drug treatment; *Right*: Centroid distances measured from indicated drug concentration to p.M1879T vehicle control. Data in the PCA plots is from Run 1.

## Discussion

In this work we performed deep electrophysiological phenotyping of human iPSC-derived neurons with the *SCN2A* p.M1879T variant, a gain-of-function pathogenic variant associated with infantile-onset epilepsy and later pervasive neurodevelopmental delay. By using a high throughput all-optical electrophysiology platform, we were able to assay the intrinsic excitability properties of >30,000 individual neurons at a single time point, a feat impossible using standard patch-clamp recordings. These measurements showed that p.M1879T patient-derived neurons have higher and longer firing than isogenic control neurons, indicating a reduced propensity for depolarization block and failure of firing. This phenotypic separation was also observed when comparing less intense optogenetic stimuli with longer stimulus duration.

Our complementary data with manual patch-clamp electrophysiology found higher action potential amplitude and accelerated upstroke of the action potential in variant neurons. Taken together, these findings support the pathogenicity of channel gain-of-function inferred from biophysical characterization of this variant in heterologous cells. In both manual patch clamp^25^ and high-throughput electrophysiology^22^, the p.M1879T variant exhibited a depolarizing shift in voltage-dependent inactivation and slower fast inactivation kinetics, leading to greater channel availability.

The neuronal phenotype we defined in this work is strikingly similar to that of the nearby *SCN2A* p.R1882Q recurrent variant, which is implicated in developmental epileptic encephalopathy and frequently associated with medically intractable epilepsy. Using manual patch-clamp of iPSC-derived neurons, Mao et al.^18^ found that the p.R1882Q neurons displayed higher firing rates at the highest stimuli compared to the isogenic controls. This difference was seen at DIV20-22, an earlier time point than the one used in our work (DIV 32+).

To understand how clinically available sodium channel blockers are efficacious in gain-of-function *SCN2A*-related disorders, we performed pharmacologic profiling of both p.M1879T and isogenic control neurons using four sodium channel blockers with different mechanisms. All four anti-seizure drugs showed a dose-dependent reduction in stimulus-firing curves, allowing for reversal of the p.M1879T phenotype. Using the all-optical electrophysiology platform, we demonstrated some of the concentrations of different anti-seizure drugs reversed the increased firing of the p.M1879T lines, rendering them indistinguishable from the isogenic control neurons. These drug concentrations therefore might have targeted anti-seizure efficacy by restoring the aberrant firing of patient neurons, while potentially minimizing side effects that occur due to blockade of physiologic neuronal firing.

Interestingly, both lacosamide and cenobamate showed similar profiles in reversing the p.M1879T phenotype, indicating the clinical applicability of these newer agents in *SCN2A*-related disorders with gain-of-function pathophysiology. These drugs lack the drawbacks seen with more conventional sodium channel blocking drugs, with a higher therapeutic index and less drug-drug interactions compared to carbamazepine and phenytoin^32^. Cenobamate has shown efficacy in medically intractable epilepsy as an adjunct therapy. It has a postulated dual mechanism for efficacy in seizure prevention: blockade of sodium channel non-inactivating persistent current, and agonism of GABA-A receptors^31,33^. The anti-seizure mechanism for lacosamide is thought to be mediated at least partially through enhancement of the slow inactivation of voltage-gated sodium channels^30,34,35^. Similar to our study, treatment of the gain-of-function p.R1882Q neurons with phenytoin reduced firing frequencies close to control levels^18^. Our work expands upon this finding in gain-of-function neurons by employing several clinically available sodium channel blockers, taking advantage of the high-throughput all-optical electrophysiology approach.

Both the p.M1879T and p.R1882Q variants predominantly affect voltage-dependent inactivation in heterologous cells^22,24,25,36^. These variants have now been shown to have similar neuronal effects, with greater firing seen at higher stimuli. It is then plausible that other *SCN2A* missense variants predominately affecting inactivation properties will have the same neuronal effect, resulting in increased firing during periods of prolonged stimulation, a phenomenon which may lead to increased seizure propensity. Combining results of high-throughput patch clamp and traditional manual patch clamp in heterologous cells, there are four pathogenic variants (in addition to M1879T and R1882Q) which can be thus defined: R1626Q, S1780I, E1880K, R1882L. Interestingly, all of these variants cluster in the last domain of the sodium channel, with four of them in the proximal C-terminus (<u>M_1879_E_1880_</u>E<u>R_1882_)</u>, with their respective locations shown in **Figure 6**. A recent study^37^ demonstrates the translational potential of an engineered peptide (ELIXIR) for sodium channel variants with aberrant inactivation, and thus this peptide may provide a promising therapy for patients with pathogenic variants in this location due to their shared biophysical mechanism.

**Figure 6.**
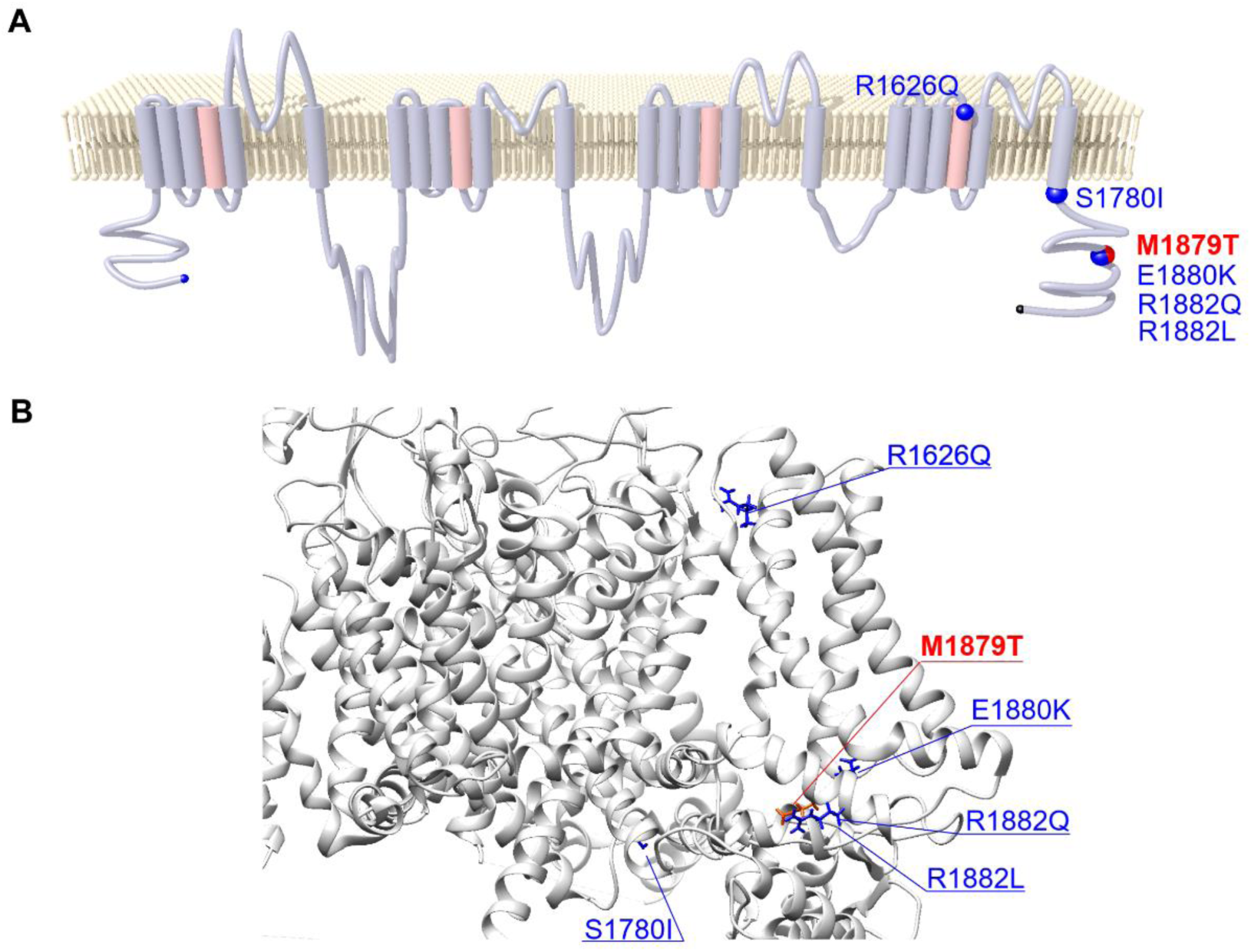
Location of Variants in NaV1.2 Predominantly Affecting Voltage-Dependent Inactivation A. Two dimensional topological structure of Nav1.2 showing location of missense variants highlighted in the text (M1879T in red; others in blue). B. Three dimensional structure with these same missense variants; structure based on PDB 6J8E (transmembrane) and 4JPZ (C-terminal domain).

Our study indicates that the *SCN2A* p.M1879T variant, associated with early-onset epilepsy and later pervasive neurodevelopmental delay, results in aberrant neuronal firing especially at higher and longer stimuli. The human iPSC-derived neurons expressing this variant continue to fire longer into the stimulus, consistent with less of the depolarization block ‘brake’ occurring in healthy neurons. We used clinically available sodium channel blocker anti-seizure drugs to reverse the firing phenotype, showing that specific drug concentrations could correct the *in vitro* functional abnormalities detected in the human neuronal model of SCN2A-related disorders.

Our findings provide new mechanistic insight into how a gain-of-function pathogenic variant primarily affecting sodium channel fast inactivation can lead to excitatory neuron dysfunction, and how targeted pharmacological treatment with drugs used in the clinic can reverse this dysfunction.

## Materials and Methods

### Study Participant

The patient was enrolled in study #2015-738, approved by the Institutional Review Board of Ann & Robert H. Lurie Children’s Hospital of Chicago. Following informed consent, data were collected by parental report through a REDCap database questionnaire and review of medical records.

### Generation and culture of induced pluripotent stem cell (iPSC) lines

The p.M1879T patient iPSC line was generated in a manner similar to what was previously described^29^. Briefly, peripheral blood mononuclear cells (PBMCs) were collected from the proband with *SCN2A* p.M1879T variant. These cells were then transduced with Sendai virus for reprogramming using Yamanaka factors (Oct4, Sox2, Klf4, c-Myc) and selected with dilution cloning. Pluripotency was confirmed with SSEA4 and Nanog immunocytochemistry staining.

The cell lines were sent for G-band karyotyping to confirm normal chromosomal integrity (Cell Line Genetics, Madison WI). For iPSC culture and expansion, cells were cultured on Matrigel^TM^ (Corning) and maintained in mTeSR^TM^ Plus medium (Stem Cell Technologies) which was exchanged every other day. When confluent, iPSC lines were passaged with ReLeSR^TM^ (Stem Cell Technologies). For routine passage, 10 μM Rock inhibitor (Y27632, Stem Cell Technologies) was added to the media for 24 hours.

### Generation of Isogenic-corrected iPSC lines

We used CRISPR/Cas9-mediated gene editing to correct the *SCN2A* p.M1879T variant in the patient-derived iPSC line and generate an isogenic set of lines. Guide RNAs (custom crRNA and trRNA) targeting the relevant *SCN2A* locus were assembled according to manufacturer’s instructions (Integrated DNA Technologies, IDT). The gRNA was combined with recombinant Cas9 protein (IDT) and a custom homology-directed repair (HDR) single stranded oligonucleotide (synthesized by IDT) containing the corrected *SCN2A* coding sequence along with a silent PAM site mutation (**Supplemental Table 1**). The Cas9/gRNA complex was electroporated into Accutase-dissociated iPSCs (StemCell) using the MaxCyte system (MaxCyte). After 48 hours of recovery, iPSCs were passaged and the incorporation of the *SCN2A* edit was confirmed with PCR followed by Sanger sequencing. Cells were then treated with Accutase, and single iPSCs were plated using clonal dilution onto 10 cm Matrigel^TM^-coated plates, cultured in mTeSR^TM^ Plus supplemented with CloneR^TM^ (Stem Cell Technologies) for four days and then switched to mTeSR^TM^ Plus media. After several days in culture, individual iPSC colonies were manually picked under a microscope and expanded. Two isogenic iPSC clones were isolated (Clone A and Clone B, ‘Isogenics’), along with two clones that underwent the same procedure but retained the *SCN2A* p.M1879T variant (clone C and clone D, ‘p.M1879T lines’). Sanger sequencing of PCR amplicons containing the *SCN2A* variant region confirmed the genotypes of all clones (ACGT Inc). Potential off-target sites for *SCN1A* and *SCN3A* were also inspected with Sanger sequencing of targeted PCR amplicons. The presence of noncoding SNVs in the isogenic clones confirmed heterozygosity.

### Neuronal Differentiation and Culture

Cortical excitatory neurons were produced using NEUROG2 (NGN2)-mediated transcriptional programming^28^ combined with forebrain patterning molecular cues (dual SMAD/Wnt inhibition). Accutase-dissociated iPSCs were transduced with lentiviral particles (produced as described previously^27^) encoding the reverse tetracycline transactivator [FUdeltaGW-rtTA (addgene #19780)] and doxycycline-responsive NEUROG2 [tetON-hNGN2-P2A-EGFP-Puro (addgene #79823) and plated at 100,000 cells/cm^2^ in mTeSR^TM^1 supplemented with Rock Inhibitor (Y-27632, StemCell). On day 1 of differentiation, cell culture medium was changed to Knockout Serum Replacement (KSR, Gibco) medium supplemented with 10 μM SB431542 (StemCell), 2 μM XAV939 (StemCell), and 100 nM LDN-193189 (StemCell) along with doxycycline (Sigma, 3 μg/mL). Day 2 medium was 1:1 KSR and Neural Induction Media (NIM) supplemented with SB/XAV/LDN/dox and puromycin (2 μg/mL). On Day 3, medium was exchanged to NIM with puromycin and doxycycline. On Day 4, differentiated cells were dissociated with Accutase and then cryopreserved in 10% DMSO (Sigma) and a 1:1 mix of fetal bovine serum (FBS, R&D Systems) and Neurobasal supplemented with B27/N2 (Gibco) and 10 ng/mL brain-derived neurotrophic factor (BDNF, R&D Systems). For functional assays, differentiated neurons were thawed and resuspended in Neurobasal medium supplemented with N2, B27 and Rock Inhibitor and plated on pre-coated Poly-D-lysine/laminin coverslips containing a supportive monolayer of rodent glial cells (seeded 4-5 days previously, isolated as described previously^29^). The differentiated neurons were kept in Neurobasal medium supplemented with N2, B27, BDNF, and doxycycline (2μg/mL) for the remainder of culture. The day after plating, differentiated cells were treated with 4 μM AraC (Tocris) for 3 days to eliminate undifferentiated dividing cells. For all-optical electrophysiology experiments, a nearly identical protocol was used except for EdU (5-ethynyl-2’-deoxyuridine, Thermo Fisher Scientific) treatment in place of AraC.

### Whole-cell electrophysiology

Coverslips with cultured neurons were transferred to a chamber continuously perfused with room temperature artificial CSF and bubbled with 5% CO2 carbogen. Recordings were sampled at 50 kHz and lowpass filtered at 10 kHz. Recordings were digitized with Digidata 1550B and acquired with Multiclamp 700B headstage using PClamp software (Molecular Devices). Pipette resistances were 3-5 MegaOhms. External solution aCSF contained (in mM): 124 NaCl, 4.4 KCl, 2.4 CaCl_2_, 1.3 MgSO_4_, 1 NaH_2_PO_4_, 26 NaHCO_3_, Glucose 10, HEPES, pH 7.35, bubbled with carbogen. Pipette solution contained (in mM): 110 K-Gluconate, 10 KCl, 10 Phosphocreatine-Tris, 5 EGTA, 10 HEPES, 4 Mg-ATP, 0.3 Na-GTP, 0.1 CaCl_2_, 10 NaCl, pH 7.35. Access resistance was monitored throughout the recording; cells with >25 MegaOhms were discarded from further analysis. Cells were recorded in both current clamp and voltage clamp. Analysis was performed with Clampfit (Molecular Devices). All electrophysiology data was from at least three independent differentiations per *SCN2A* genotype.

### All-Optical electrophysiology (*Optopatch*)

Neuronal culture for *Optopatch* measurements was carried out as previously described with some minor modifications^38^. Cryostocks of differentiated neurons were thawed and plated at a density of 50,000 cells per well onto custom-made 96-well Ibidi^TM^ cell culture plates containing a monolayer of primary mouse glial cells (seeded 7 days before). Co-cultures were maintained in Neurobasal medium supplemented with N2, B27, and neurotrophic factors BDNF and GDNF. Neurons were transduced on day *in vitro* 21 (DIV21) with lentiviral particles encoding all-optical physiology components CheRiff-BFP (channelrhodopsin) and QuasAr3-Cirine (voltage indicator), with targeted expression to mature neurons by means of the human SYNAPSIN 1 (*hSYN1*) promoter. Functional experiments were performed two weeks after transduction on DIV35. After primary data acquisition, segmentation, source de-mixing, and functional feature extraction of voltage imaging movies were conducted as previously described^27,38^. Two independent runs of neuronal culture and optical physiology measurements were performed.

For pharmacology experiments, culture wells were treated with selected drugs at different concentrations (cenobamate, phenytoin, carbamazepine, and lacosamide) for 48 hours prior to stimulation and functional measurements, and neurons were recorded in the same concentration of freshly applied drug or DMSO vehicle control in imaging buffer. Extracted functional parameters from the evoked firing protocol were analyzed with custom Python scripts.

### Gene Expression Assays

Total RNA was purified from DIV 55 neurons grown on 12mm coverslips grown on coverslips from each genotype using the RNA extraction kit from Takara Biosciences according to the manufacturer’s instructions. Briefly, media was aspirated and cells were collected in RA1 solution supplemented with beta-Mercaptoethanol. RNA was column purified and contaminating genomic DNA was digested with recombinant DNAse. Approximately 100ng of RNA was reverse transcribed with the Superscript IV kit using random hexamers. The cDNA was run in a ddPCR reaction (BioRad) with primers targeting human *SCN2A* and *SCN3A* (Hs00221379_m1, Hs00366902_m1; Thermo Fisher Scientific). TATA-binding protein (TBP, Hs99999910_m1) was used as a control for normalization. For the allele-specific reaction, custom primers were ordered from Thermo Fisher Scientific with FAM (variant) and VIC (wildtype) labels.

### Machine Learning and Classification

Parameters from individual neurons were included if they met initial filtering criteria of an average signal-to-noise ratio (SNR) greater than 6.0 and at least two total spikes during the recording protocol. A bootstrap protocol was implemented for feature selection and model training using batch sizes of 100 neurons per genotype, with feature selection and training performed using the batch average over 50 iterations. Embedded feature selection was performed using a random forest classifier, with the resulting feature set fixed and applied unchanged to all test data. Classification accuracy was evaluated for each condition using independent testing groups that were not included during feature selection or model training. For aggregate analysis and PCA, neuron features were averaged at the well level, then elastic net regression was applied to remove highly correlated features. Principal components were computed to account for 95% of variability. The Euclidean distance was measured between calculated centroids.

### Statistics

Statistical testing was performed with Graphpad Prism. Datasets were assessed for normality and either a parametric (t-test) or nonparametric (Mann-Whitney, Kruskal-Wallis) test was used as indicated. Multiple comparisons were corrected using Dunn’s posthoc testing.

## Acknowledgments

We wish to acknowledge the patient and family for involvement in this research and the *SCN2A* community as a whole. We thank Dina Simkin, Ph.D., for her assistance and guidance in methodology for this project.

## Funding

This work was supported by NINDS grants U54NS108874 (ALG) and K08NS121601 (SKA)

## Contributions

Conceptualization: LAW, OM, GTD, ALG, SKA

Funding acquisition: SKA, ALG

Data curation: SJR, SP, SKA

Formal analysis: SP, DJIR, SJR, SKA

Investigation: DJIR, LAW, ST, COO, LL, ER, SKA

Methodology: SP, DJIR, LAW, ST, KH, VJ, SJR, COO, OM, GTD, SKA

Project administration: SKA

Supervision: OM, GTD, SKA

Visualization: SP, DJIR, SJR, SKA

Writing – original draft: SKA

Writing – review & editing: SP, DJIR, LAW, SJR, ALG, SKA

---

**Supplementary Table 1:**
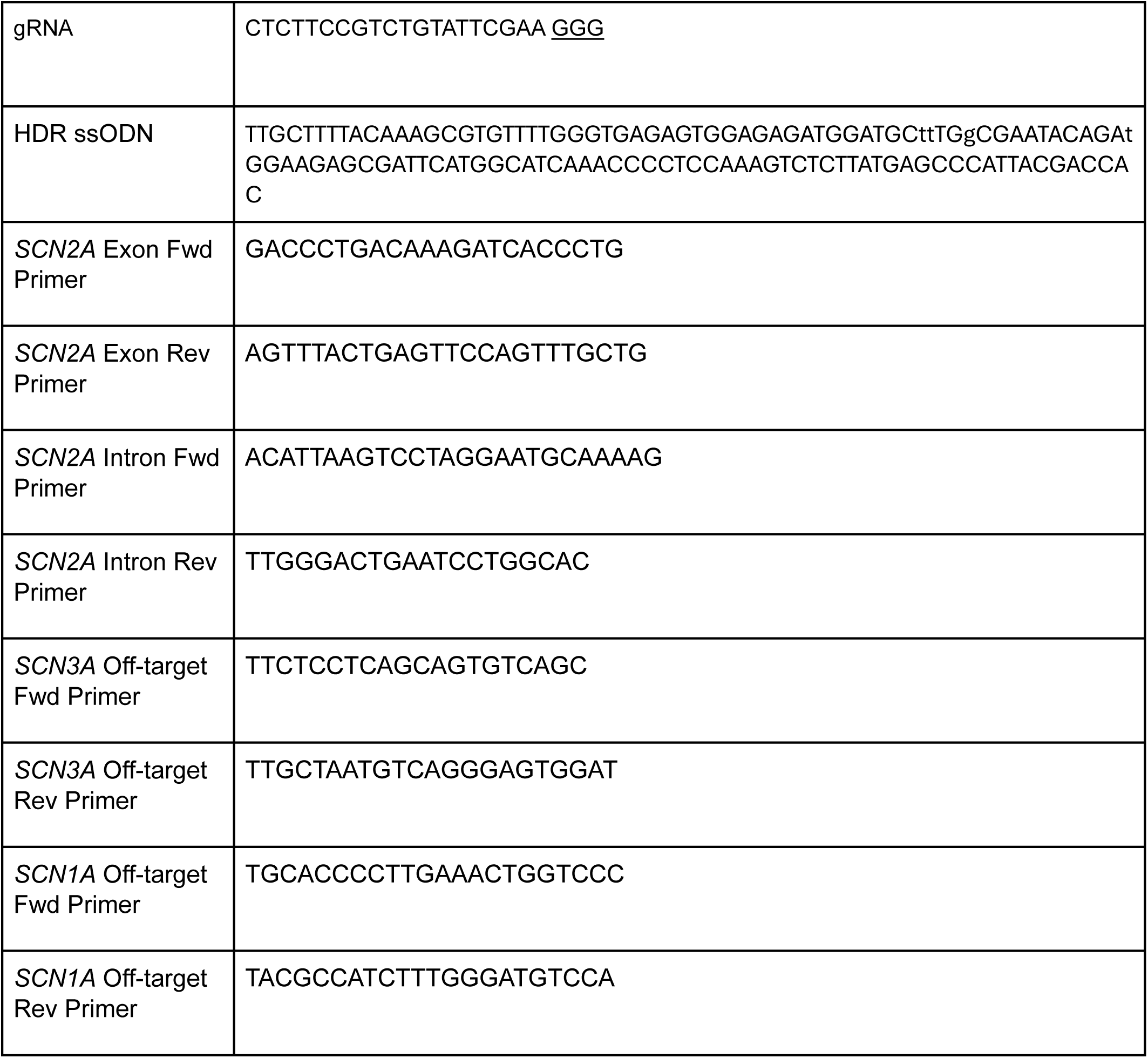
Nucleotides.

**Supplementary Figure 1.**
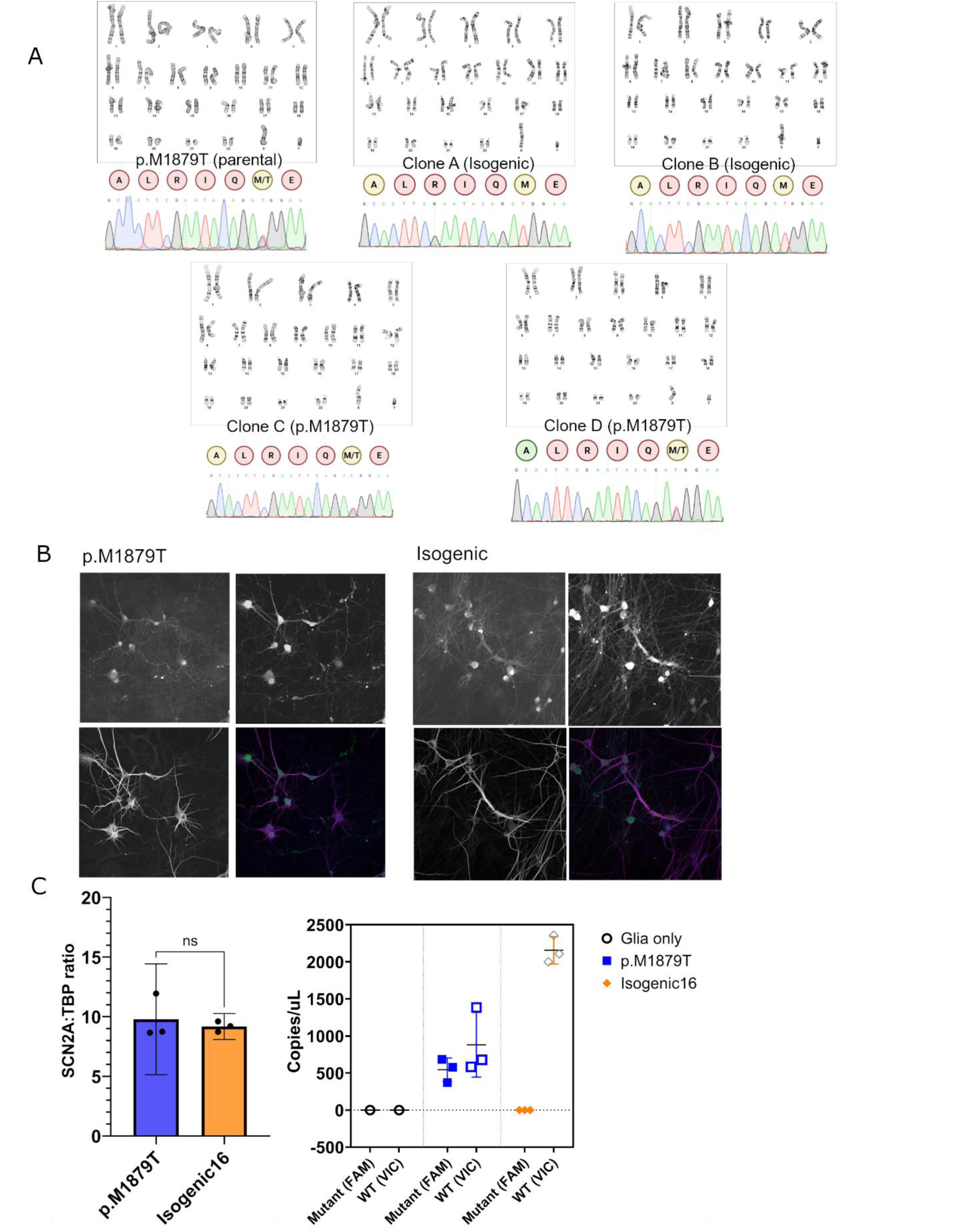
A. Karyotype and representative Sanger chromatograms for each clonal line. B. Expression of vGlut (upper left) and MAP2 (bottom left) markers in iPSC-derived neurons from two representative lines (patient and isogenic). EGFP shown in upper right panel and merged signals are on bottom right panel. C. *Left*, *SCN2A* RNA expression normalized to TATA-binding protein (TBP) with ddPCR from DIV50 neurons. *Right*, allele-specific ddPCR showing lack of mutant allele expression in isogenic control line.

**Supplementary Figure 2.**
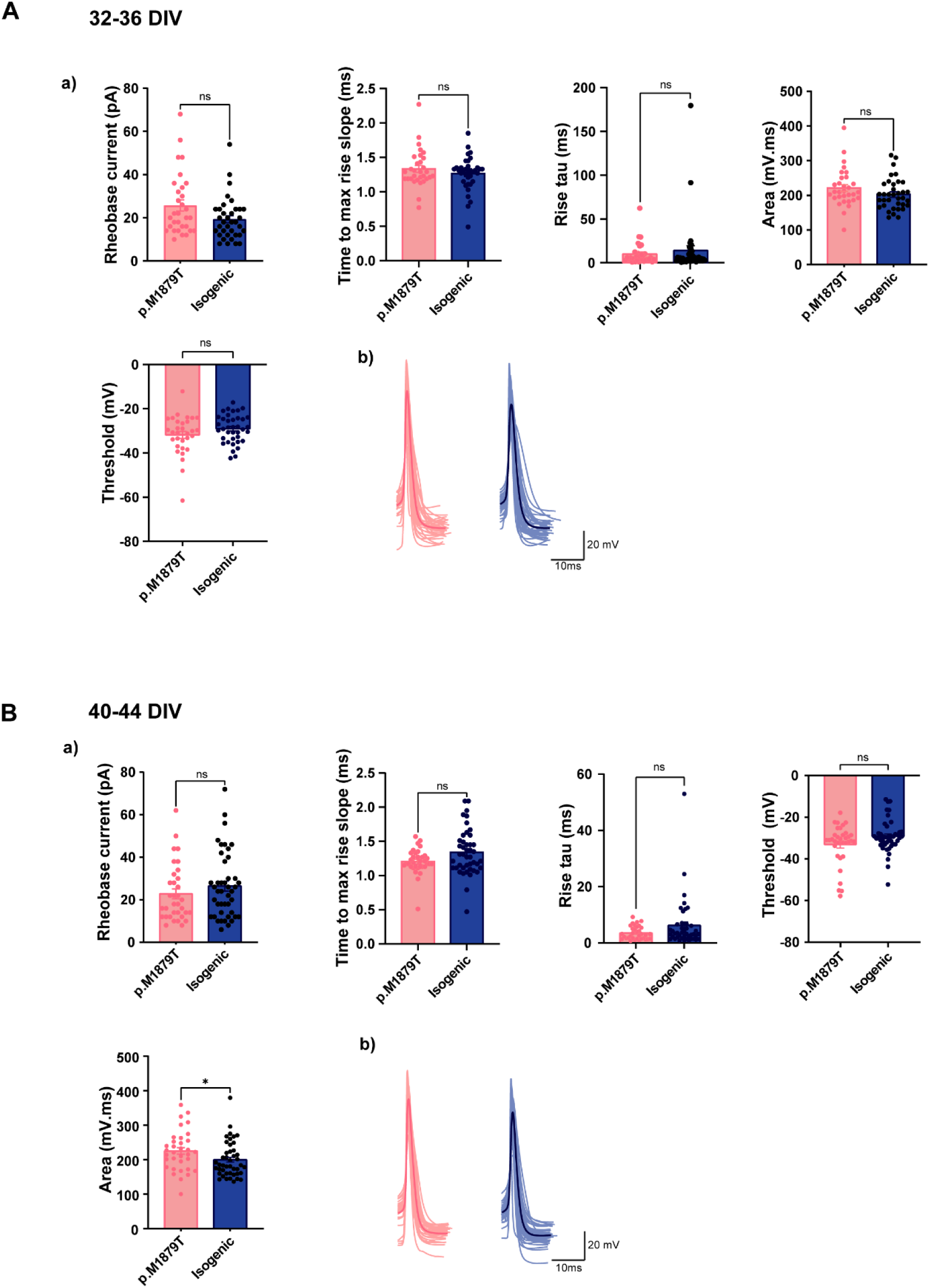
Additional electrophysiology parameters. A, B. Left to right, *top*: Summary data of rheobase, time to max rise slope, rise tau, and area of action potential. *Bottom*, *left*: Summary data of action potential threshold. *Bottom, right*: all phase plots from first evoked action potential of each neuron for p.M1879T (left) and isogenic (right). Data are from DIV32-36 (**A**) and DIV40-44 (**B**). ns, no significance by Mann-Whitney testing.

**Supplementary Figure 3.**
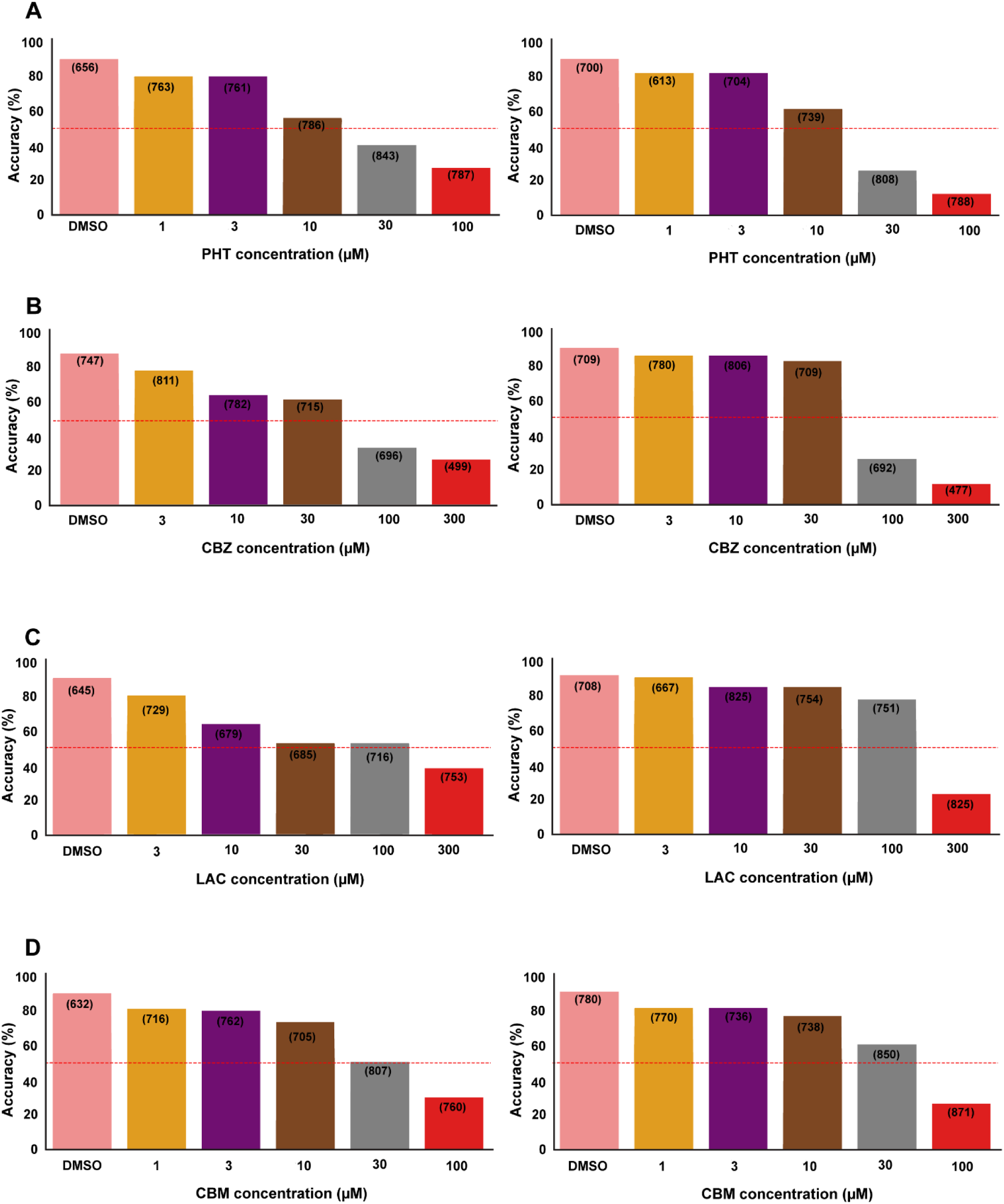
Machine learning Classification of Neurons A-D. Run-specific classification accuracy of genotype prediction of p.M1879T neurons treated with indicated drug. *Left*, Run 1 and *Right*, Run 2.

**Supplementary Figure 4.**
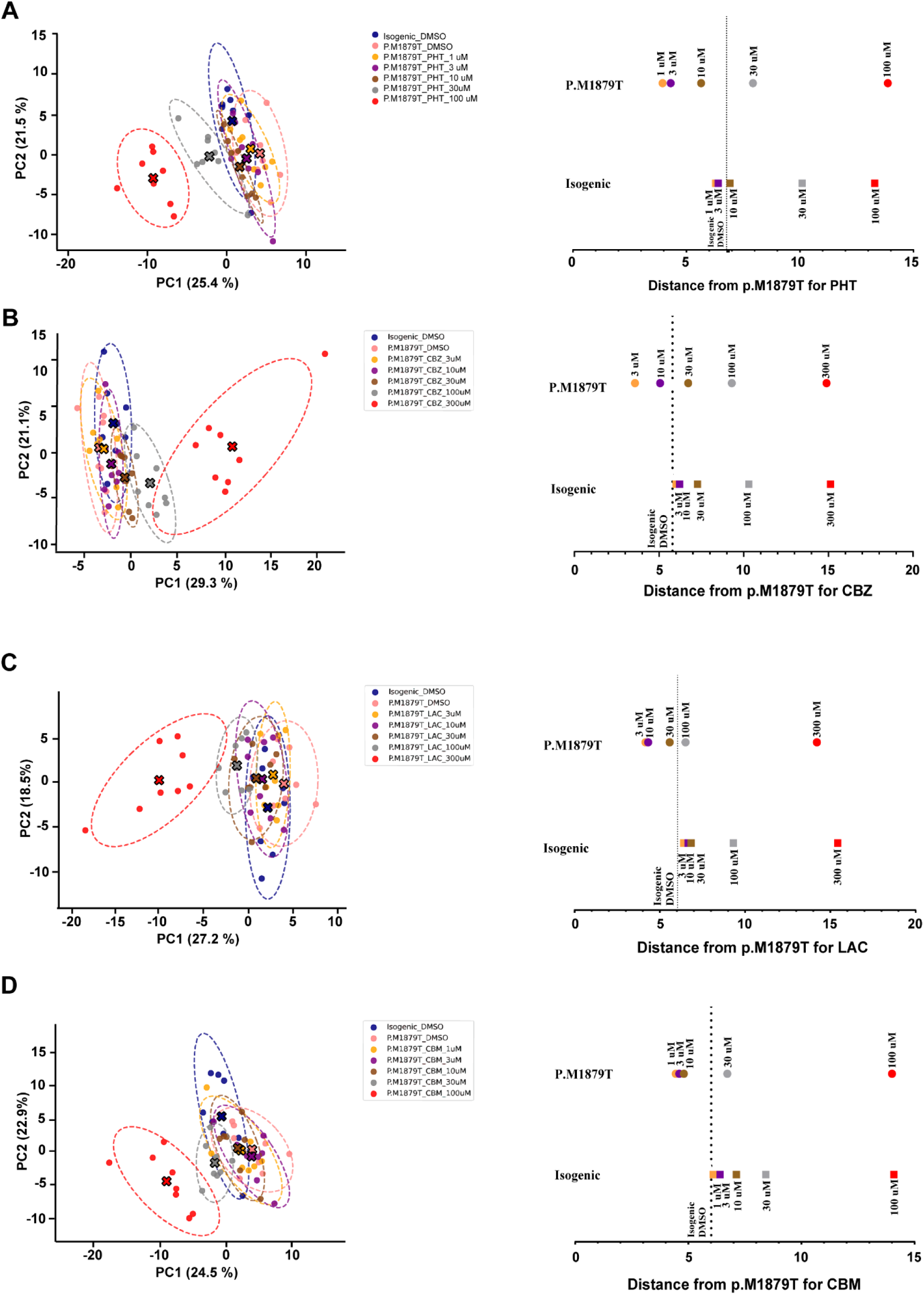
PCA analysis of Run 2 A-D. *Left*, PCA plots showing aggregate feature clusters with respect to drug treatment for Run 2; *Right*: Centroid distances measured from indicated drug concentration to p.M1879T vehicle control for Run 2.

## Notes

### Competing Interest Statement

GTD, LAW, SJR, KH, are current employees of Quiver Bioscience and hold stock options in Quiver Bioscience.
VJ and OM were employees of Quiver Bioscience during execution of the study and received stock options in Quiver Bioscience
Stock Ownership relevant to financial competing interests:
GTD, LAW, and SJR are current employees of Quiver Bioscience and hold stock options with potential value exceeding $5,000.

